# Large CRISPR-Cas-induced deletions in the oxamniquine resistance locus of the human parasite *Schistosoma mansoni*

**DOI:** 10.1101/2020.05.03.074831

**Authors:** Geetha Sankaranarayanan, Avril Coghlan, Patrick Driguez, Magda E. Lotkowska, Mandy Sanders, Nancy Holroyd, Alan Tracey, Matthew Berriman, Gabriel Rinaldi

## Abstract

At least 250 million people worldwide suffer from schistosomiasis, caused by *Schistosoma* worms. Genome sequences for several *Schistosoma* species are available, including a high-quality annotated reference for *Schistosoma mansoni*. There is a pressing need to develop a reliable functional toolkit to translate these data into new biological insights and targets for intervention. CRISPR-Cas9 was recently demonstrated for the first time in *S. mansoni*, to produce somatic mutations in the *omega-1* (*ω1*) gene. Here, we employed CRISPR-Cas9 to introduce somatic mutations in a second gene, *SULT-OR*, a sulfotransferase expressed in the parasitic stages of *S. mansoni*, in which mutations confer resistance to the drug oxamniquine. A 262-bp PCR product spanning the region targeted by the gRNA against *SULT-OR* was amplified, and mutations identified in it by high-throughput sequencing. We found that 0.3-2.0% of aligned reads from CRISPR-Cas9-treated adult worms showed deletions spanning the predicted Cas9 cut site, compared to 0.1-0.2% for sporocysts, while deletions were extremely rare in eggs. The most common deletion observed in adults and sporocysts was a 34 bp-deletion directly upstream of the predicted cut site, but rarer deletions reaching as far as 102 bp upstream of the cut site were also detected. The CRISPR-Cas9-induced deletions, if homozygous, are predicted to cause resistance to oxamniquine by producing frameshifts, ablating *SULT-OR* transcription, or leading to mRNA degradation *via* the nonsense-mediated mRNA decay pathway. However, no *SULT-OR* knock down at the mRNA level was observed, presumably because the cells in which CRISPR-Cas9 did induce mutations represented a small fraction of all cells expressing *SULT-OR*. Further optimisation of CRISPR-Cas protocols for different developmental stages and particular cell types, including germline cells, will contribute to the generation of a homozygous knock-out in any gene of interest, and in particular the *SULT-OR* gene to derive an oxamniquine-resistant stable transgenic line.

## 1. Introduction

Schistosomiasis is a major Neglected Tropical Disease (NTD) affecting more than 250 million people worldwide [1]. *S. mansoni* and *S. japonicum* are the agents of hepato-intestinal schistosomiasis manifested by abdominal pain, liver inflammation and fibrosis that leads to portal hypertension. Infection with *S. haematobium,* agent of urogenital schistosomiasis, is associated with infertility, haematuria, kidney pathology and squamous cell carcinoma of the bladder. In addition, all forms of schistosomiasis are associated with systemic morbidities that include malnutrition, anaemia, physical and/or cognitive impairment and stunted development in children [2]. Currently, praziquantel is the single effective drug to treat the infection, and is employed in mass drug administration programmes across endemic areas, which could eventually lead to drug resistance emerging [3]. Therefore, there is an urgent need for the development of novel drugs and vaccines [4]. Understanding the basic biology of schistosomes at the cellular and molecular levels is critical to identifying exploitable vulnerabilities of the parasite. High-throughput datasets, including high quality reference genomes for the three main species of schistosomes [5–7], have been generated. More recently, a thorough transcriptome analysis during the parasite’s intra-mammalian development [8], and the identification of different cell types by single-cell RNA sequencing of various life cycle stages [9, 10] represent significant steps towards deciphering cell fate and pathways involved in parasite development and host-parasite interactions.

In parallel to the creation of large-scale datasets, a functional genomics toolkit is needed to experimentally investigate hypotheses that emerge from these data, to confirm biological insights and validate targets for intervention. Recently, a large RNAi-based gene silencing screen, encompassing almost one third of *S. mansoni* protein-coding genes, revealed genes associated with parasite viability and potential targets for drug development [11]. However, not every gene is susceptible to RNAi, the effect is transient and highly variable depending on the expression level of the target gene, tissue localisation and half-life of the mRNA and protein. In addition, off-target effects are common, in particular when long dsRNA molecules are used, and the gene silencing is typically not heritable unless a RNAi-based construct is employed as a transgene expressed in the germ line [12]. Therefore, to truly examine gene function across the life-cycle, transgenesis-based approaches that are available in model organisms [13, 14] need to be developed for *S. mansoni,* including the ability to create genetically-modified parasite strains with homozygous gene knock-outs, and specific gene mutations.

Some progress with transgenesis and genome editing has been made. Retrovirus transduction of schistosome developmental stages, including eggs, has proved effective, and will likely be a key delivery system in the generation of stable transgenic lines [15–18]. Site-specific integration of transgenes and highly precise site-specific genome editing using CRISPR-Cas technology will be a key step [19]. CRISPR-Cas9 has recently been used to create a heritable gene knock-out line in the parasitic nematode *Strongyloides stercoralis* [20]. In *S*. *mansoni*, the technology has been used to produce mutations in the *omega-1* (*ω1)* gene in somatic cells of the egg [21], and in a related parasitic flatworm, the liver fluke *Opisthorchis viverrini*, CRISPR-Cas9 mutations have been introduced into the granulin gene in somatic cells of adult worms [22]. Somatic mutations in these two flatworms were associated with dramatic reductions in *ω1* and granulin mRNA levels, respectively, and produced *in vitro* and *in vivo* phenotypic effects shedding new light on their functional roles and contributions to pathogenesis [21,22].

Whether different CRISPR-Cas protocols are needed to deliver site-specific mutations in *S. mansoni* genes expressed in other tissues or developmental stages remains to be determined. Likewise, the types of mutations to be expected and the degree of mRNA knock-down in different genes is not yet known. In the current study we have used CRISPR-Cas9 to introduce site-specific mutations in a second *S. mansoni* gene, to better understand how the CRISPR-Cas system works when applied to *S. mansoni.* We compared the efficiency of the approach in different developmental stages of the parasite: eggs, mother sporocysts (the first intra-snail stage) and adult worms. The mutations produced by CRISPR-Cas9, including their sizes and locations, were characterised. We chose the *SULT-OR* sulfotransferase (*Smp_089320*) gene as a target because recessive mutations in this gene, both induced in laboratory conditions and detected in field samples, confer resistance to the drug oxamniquine (OXA) [23, 24]. In addition, it is mostly expressed in the intra-mammalian stages of the life cycle (schistosomula and adults) [25] that would likely be the target of any new intervention strategy. Our findings provide insights that will help pave the way towards using CRISPR-Cas to achieve the generation of stable genetically-engineered schistosomes.

## 2. Materials and methods

### 2.1 Ethics statement

The complete life cycle of *Schistosoma mansoni* NMRI (Puerto Rican) strain is maintained at the Wellcome Sanger Institute (WSI) by breeding and infecting susceptible *Biomphalaria glabrata* snails and mice. The mouse experimental infections and rest of regulated procedures were conducted under the Home Office Project Licence No. P77E8A062 held by GR. All protocols were revised and approved by the Animal Welfare and Ethical Review Body (AWERB) of the WSI. The AWERB is constituted as required by the UK Animals (Scientific Procedures) Act 1986 Amendment Regulations 2012.

### 2.2 Parasite material

Developmental stages of *S. mansoni* were collected and maintained as described [26]. In brief, mixed-sex adult worms were collected by portal perfusion of experimentally-infected mice 6 weeks after infection, washed with 1x Phosphate-Buffered Saline (PBS) supplemented with 200 U/ml penicillin, 200 μg/ml streptomycin and 500 ng/ml amphotericin B, and cultured in complete DMEM medium (high-glucose DMEM, 10% Fetal Bovine Serum (FBS), 200 U/ml penicillin, 200 μg/ml streptomycin and 500 ng/ml amphotericin B; all media components were purchased from ThermoFisher Scientific) at 37°C, under 5% CO_2_ in air. *S. mansoni* eggs were isolated from the livers of experimentally-infected mice removed after the portal perfusion [27]. The livers were finely minced and digested overnight in the presence of 0.5% *Clostridium histolyticum* collagenase (Sigma), followed by three washes with 1x PBS and filtered through 250 μm and 150 μm sieves. The filtrate was passed through a Percoll-sucrose gradient, and the resulting purified eggs washed in 1x PBS and cultured in complete DMEM medium at 37°C, under 5% CO_2_ in air as described [28]. Primary sporocysts were obtained by transferring miracidia hatched from freshly collected eggs into complete sporocyst medium (MEMSE-J, 10% Fetal Bovine Serum, 10mM Hepes, 100 U/ml penicillin, 100 μg/ml streptomycin) and cultured in a hypoxia chamber in a gas mixture of 1% O_2_, 3% CO_2_ and balance N_2_, at 28°C [26].

### 2.3 CRISPR-Cas 9 ribonucleoprotein complex assembly

We explored the activity of a ‘two-piece’ guide RNA that included a (1) CRISPR RNA (crRNA) molecule of 20 nucleotides target-specific sequence, and (2) the conserved 67 nucleotide trans-activating crRNA (tracrRNA). The crRNA sequence 5’-ACAATCCAAGTTATCTCAGC-3’, spanning positions 19-38 from the first codon of exon 1 of *SULT-OR* (*Smp_089320*) and followed by the protospacer adjacent motif (PAM) TGG **(Figure 1)**, was designed using the web-based tool CRISPR RGEN Tools (Computational tools and libraries for RNA-guided endonucleases, RGENs; http://www.rgenome.net/). The crRNA, the fluorescently labelled tracrRNA (Alt-R® CRISPR-Cas9 tracrRNA, ATTO™ 550), and the recombinant *Streptococcus pyogenes* Cas9 nuclease containing a nuclear localization sequence (Alt-R® S.p. Cas9 Nuclease V3) were purchased from IDT. The CRISPR-Cas9 ribonucleoprotein complex (RNP) was assembled *in vitro* by combining the ‘two-piece’ gRNA with the Cas9 nuclease (163.7 kDa). Briefly, the ‘two-piece’ gRNA was generated by mixing equal volumes of 200 μM *SULT-OR* crRNA and 200 μM ATTO™ 550 tracrRNA in IDT buffer. The RNA oligos were annealed by incubating the mixture at 95°C for 5 min followed by a slow cooling to room temperature for at least 10 min. Thereafter, the RNP was assembled by combining 100 pmol Cas9 nuclease (stock concentration, 10 μg/μl = 61 μM) with 150 pmol ‘two-piece’ gRNA. The RNP was gently mixed avoiding pipetting, incubated at room temperature for 10 min and kept on ice. Immediately before the parasite transfection Opti-MEM media (ThermoFisher Scientific) was added to the RNP to reach a final volume of 100 μl and kept on ice.

**Figure 1.**
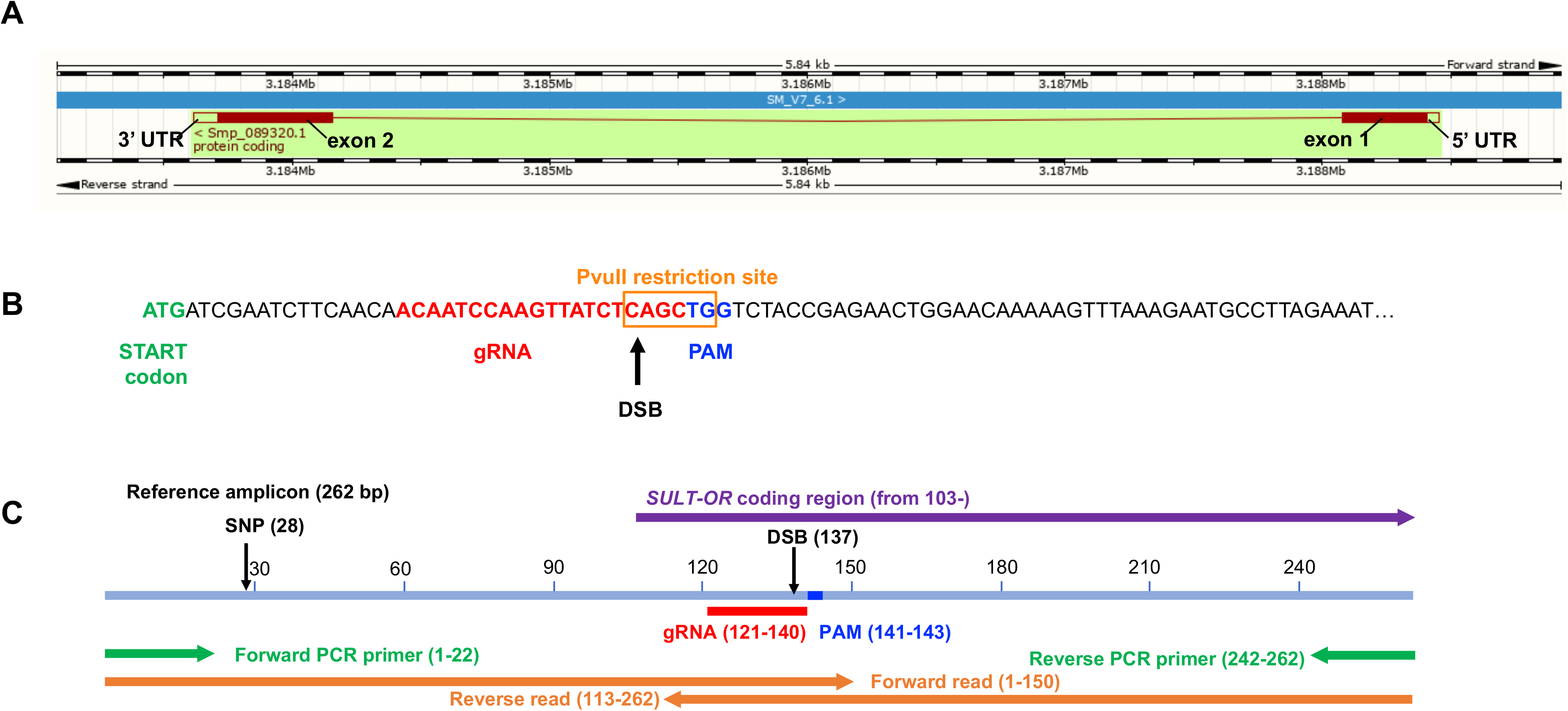
**(A)** Gene model of *SULT-OR* (*Smp_089320*), indicating the position of the two exons, one intron, and UTRs, spanning 4837 bp on the reverse strand of chromosome 6. Boxes filled in dark red represent the protein coding sequence. *Schistosoma mansoni* (PRJEA36577). Assembly: Smansoni_v7 (GCA_000237925.3). Region: Scaffold SM_V7_6:3,183,084-3,188,924. Adapted from WormBase ParaSite 14 [34]. **(B)** Nucleotide sequence of the start of exon 1 indicating location and sequence of gRNA target site, predicted double-stranded break (DSB), protospacer adjacent motif (PAM), and the PvuII restriction site. **(C)** Reference PCR amplicon, showing the positions of the gRNA, PAM, DSB, forward and reverse PCR primers, and forward and reverse sequence reads, as well as a SNP site found in many sequences reads. The diagrams are drawn to scale.

### 2.4 CRISPR-Cas9 transfection of schistosome developmental stages

The CRISPR-Cas9 RNP was delivered into *S. mansoni* mixed-sex adult worms, eggs and *in vitro*-transformed mother sporocysts by square-wave electroporation as previously described [29, 30] with minor modifications. Briefly, groups of ~16 male and female worms were transferred to a pre-cooled 4-mm electroporation cuvette (BTX), and washed 3 times by gravity with Opti-MEM medium with no FBS and no antibiotic/antimycotic mix. After the last wash, the worms were maintained in 50 μl of Opti-MEM medium and the RNP in 100 μl of Opti-MEM medium (above) was added to the cuvette containing the worms. The eggs isolated from the livers and cultured as described above were collected and washed in Opti-MEM medium (no FBS and no antibiotic/antimycotic mix) 3 times by centrifugation at 400 g for 5 min. After the last wash, the eggs were split into groups of ~10,000, resuspended in 100 μl of Opti-MEM medium containing the RNP (above) plus 50 μl of Opti-MEM to collect all the remaining eggs from the original tube, and transferred to a pre-cooled 4-mm electroporation cuvette (BTX). Three-day old *in vitro*-transformed sporocysts were collected and washed in Opti-MEM medium (no FBS and no antibiotic/antimycotic mix) 3 times by centrifugation at 400 g for 5 min. After the last wash, the sporocysts were split in groups of ~10,000, resuspended in 100 μl of Opti-MEM medium containing the RNP (above) plus 50 μl of Opti-MEM to collect all the remaining sporocysts from the original tube, and transferred to a pre-cooled 4-mm electroporation cuvette (BTX). The final electroporation volume for the schistosome worms, eggs and sporocysts was 150 μl, i.e. the final concentration of the RNP complex in the cuvette was 1.67 μM. The three developmental stages were subjected to the same electroporation conditions; square-wave, a single pulse of 125V for 20 msec in a BTX Gemini X2 electroporator (BTX). Immediately after electroporation, the schistosome worms and eggs were collected in pre-warmed complete DMEM medium, and the sporocysts in complete sporocysts medium and cultured as described above. Four hours after transfection, three male and female worms and a few thousand eggs and sporocysts were collected for confocal microscopy (below). Four days post-transfection the parasites were collected, washed in 1x PBS and processed for DNA and RNA isolation (below). In addition to the CRISPR-Cas9 experimental condition, i.e. parasites exposed to the CRISPR-Cas9 RNP complex, we included three control groups subjected to the same electroporation protocol: (1) mock-treated group that included parasites exposed to no molecules, (2) parasites exposed to Cas9 nuclease only, and (3) parasites exposed to the ‘two-piece’ gRNA only.

**Supplementary Table S1** summarises the experimental conditions and biological replicates performed for each of the three tested developmental stages.

### 2.5 DNA isolation and amplicon sequencing libraries

A conventional phenol:chloroform:isoamyl alcohol (25:24:1) protocol was employed to isolate DNA from RNP-transfected parasites and all control groups. Briefly, wet pellets of adult worms, eggs or mother sporocysts stored at −80°C were incubated overnight in the presence of 500 μl genomic DNA lysis buffer (200 mM NaCl, 100 mM Tris-HCL pH 8.5, 50 mM EDTA pH 8, 0.5 % SDS) and 10 μl of proteinase K (20 mg/ml) at 56°C with agitation (400 rpm). Thereafter, 5 μl of 4 mg/ml of RNase A was added to the lysate and incubated at 37°C for 10 min. One volume of phenol-chloroform-isoamyl alcohol (25:24:1) was added to the sample, mixed by shaking the tube vigorously, incubated at room temperature for 5 min and centrifuged at 14,000 g at room temperature for 15 min. The aqueous top layer was transferred to a new tube and 1 volume (~200 μl) of chloroform:isoamyl alcohol (24:1) was added to the sample, mixed vigorously and centrifuged as above. The aqueous top layer containing the DNA was transferred to a new tube and precipitated with 0.1 volume of 3 M sodium acetate, 3 volumes of 95%-100% ethanol, and 2 μl of Glycoblue (Thermo Fisher Scientific) overnight at −20°C. The DNA was recovered by centrifugation at 14,000 g at 4°C for 30 min, washed with 500 μl of 70% ethanol, resuspended in pre-warmed nuclease-free water and quantified by Qubit fluorometer. For the indicated samples **(Supplementary Table S1)** in order to enrich for *SULT-OR* mutant alleles, 20 ng of DNA was digested with 6 to 12 U of the restriction enzyme PvuII (NEB) overnight at 37°C.

For the amplicon library preparation, a 2-step PCR protocol was followed. During the first PCR, a 262 bp *SULT-OR-*specific amplicon spanning the predicted double-stranded breaking site (DBS) was generated using 10 ng template DNA (20 μl of 0.5 ng/μl DNA preparation), 300 nM forward and reverse primers **(Supplementary Table S2)**, and 2x Kapa HiFi Master Mix (Roche) in a 50 μl PCR reaction performed in a Thermocycler (Eppendorf mastercycler). The PCR protocol included an initial denaturation step at 95°C for 3 min, 18 cycles of denaturation step at 98°C for 20 sec, annealing step at 53°C for 15 sec, and extension step at 72°C for 40 sec, followed by a final extension step at 72°C for 5 min. For sample DNA preparations that were digested with PvuII, two PCR reactions per sample were run in parallel and the products were pooled, i.e. a total of 20 ng of each of two PvuII-digested DNA preparations was used to generate the amplicon. Four PCR reactions per sample were run in parallel for the rest of the samples and the products were pooled at the end, i.e. a total of 40 ng of each sample DNA preparation was used to generate the amplicon. The pooled PCR products for each sample were cleaned up using a column-based kit (Zymo DNA Clean and concentrator), eluted in 17 μl of nuclease-free water; 2 μl were used for quantification and the rest entirely used as template in the second PCR for Nextera Indexing (Nextera-XT Index kit-FC-131-1001). In a 50 μl-reaction the concentrated DNA (15 μl) was mixed with 10 μl of the Nextera index mix (i5 + i7) and 2x Kapa HiFi Master Mix (Roche). The PCR was performed in a Thermocycler (Eppendorf mastercycler) with an initial denaturation step at 95°C for 3 min, 8 cycles of denaturation step at 98°C for 20 sec, annealing step at 55°C for 15 sec, and extension step at 72°C for 40 sec, followed by a final extension step at 72°C for 5 min. The PCR products were purified using a bead-based cleaner kit (AMPure XP, Beckman Coulter), eluted in 30 μl of nuclease-free water and quantified using a high sensitivity DNA chip in a Bioanalyzer (Agilent 2000). Equimolar amounts of each library were combined and 20-30% PhiX was added to the mix to introduce complexity into these low-diversity amplicon libraries.

### 2.6 Bioinformatic analysis

Amplicon libraries from the samples summarised in **Supplementary Table S1** were sequenced on a MiSeq Illumina sequencing platform spiked with 20-30% PhiX to generate diversity. If a sample had been multiplexed and run on several MiSeq lanes, the fastq files for that particular sample were merged. Trimmomatic [31] was used to discard low quality read-pairs where either read had average base quality < 23. To detect CRISPR-induced mutations, the software CRISPResso v1.0.13 [32, 33] was employed using a window size of 500 bp (-w 500) with the reference amplicon according to *Smp_089320* in the *S. mansoni* V7 assembly from WormBase ParaSite [34]. In most samples, the majority of reads had a G→A SNP at position 28 of the amplicon, presumably due to genetic variation in the population of *S. mansoni* NMRI strain in our laboratory. Thus, although the *S. mansoni* V7 reference assembly has ‘G’ at this position, we used ‘A’ at this position in the ‘reference amplicon’ sequence given to CRISPResso. A window size of 500 bp was used to include the entire amplicon. CRISPResso was run with the -exclude_bp_from_left 30 and - exclude_bp_from_right 30 options in order to disregard the (21-22 bp) primer regions on each end of the amplicon, and the SNP at position 28, when indels and substitutions were being quantified and reads being classified as ‘NHEJ’ or ‘unmodified’ by CRISPResso.

### 2.7 Gene expression analysis for *SULT-OR* gene

Total RNA was extracted from adult worms, eggs or *in vitro* transformed sporocysts following a phenol:chloroform-based protocol. In brief, four days after transfection, parasites were collected from the culture, washed three times in 1x PBS complemented with antibiotic-antimycotic as described above for each of the three developmental stages, transferred to 1ml of Trizol, incubated at room temperature for ~10 min and stored at −80°C. The parasites in Trizol were mechanically-dissociated using a bead beater homogenizer (Fast Prep-24, MP Biomedicals) using two 20-second pulses at setting four for adult worms and *in vitro* transformed sporocysts, and two 20-second pulses at setting six, after three cycles of freezing-thawing, for eggs. Thereafter, one volume of chloroform was added to the samples, mixed vigorously, centrifuged at 14,000 g at room temperature for 15 min, and the aqueous top layer was carefully transferred to a clean tube. The total RNA was precipitated using an equal volume of 100% molecular biological grade ethanol. Residual DNA was removed by digestion with DNaseI (Zymo). RNA was cleaned and concentrated using Zymo RNA clean and concentrator columns, and eluted in 15 μl of nuclease-free water. cDNA was synthesized from 65 −175 ng of total RNA using the iScript cDNA Synthesis Kit (Bio-Rad, Hercules, CA). Target-specific primers designed with the assistance of the free web-based PrimerQuest® Tool (IDT) are shown in **Supplementary Table S2**, and the amplification efficiencies for each primer set were estimated to be 90-105% by titration analysis [35]. Real time quantitative PCRs (qPCR) were performed in triplicate, in 96-well plates, following an initial denaturation step at 95°C for 3 min followed by 40 cycles of 30 sec at 95°C and 30 sec at 50 °C, and a final melting curve, in a StepOnePlus™ Real-Time PCR System (Applied Biosystems). Reactions run in 10 μl included 300 nM of each target-specific primer, 1 μl of cDNA, and Kapa Sybr FastqPCR Master Mix (Roche). The relative quantification assay [36] was employed using *S. mansoni* glyceraldehyde-3-phosphate dehydrogenase (*SmGAPDH*, *Smp_056970*), and *S. mansoni* α-tubulin1 (Sm*AT1*, *Smp_090120*), as reference genes. The target gene expression levels were normalised using the control group.

### 2.8 Confocal microscopy

Four hours after electroporation with fluorescently labelled RNP complex, transfected adult worms, eggs or mother sporocysts were collected from the culture, fixed and processed for confocal microscopy imaging. In brief, the parasites were collected and washed three times in 1x PBS complemented with antibiotic-antimycotic solution as described above; adult worms were washed by gravity, and eggs and sporocyst by centrifugation, 400 g for 5 min. After the final wash the parasites were fixed overnight in 4% methanol-free paraformaldehyde (Pierce™) diluted in 1x PBS at 4°C, washed three times in 1x PBS, resuspended in mounting media containing 4’, 6’-diamidino-2-phenylindole (DAPI) for nuclear staining (Fluoromount-G™ Mounting Medium, with DAPI, Invitrogen), and incubated overnight at 4°C. The parasites were mounted on microscope slides and images taken with a Leica SP8 confocal microscope using appropriate settings to capture DAPI and ATTO 550 fluorochromes. Manipulation of digital images was undertaken with the assistance of the LAS X software (Leica) and was limited to insertion of scale bars, adjustment of brightness and contrast, and cropping. The image enhancement algorithms were applied in a linear fashion across the entire image.

### 2.9 Accession numbers

The sequence data generated in this study are available at the European Nucleotide Archive (ENA) accession number ERP 121238. The accession numbers for each sample is shown in **Supplementary Table S1**, columns P, Q.

## 3. Results

### 3.1 The *SULT-OR* gene belongs to a multi-copy locus on chromosome 6 of *S. mansoni*

The *SULT-OR* gene (*Smp_089320*) belongs to a multi-copy locus containing six other paralogous genes on chromosome 6 of the *S. mansoni* reference genome, version 7 (WormBase ParaSite, https://parasite.wormbase.org) **(Supplementary Figure S1)**. The biological function of SULT-OR remains unknown, except that it converts the pro-drugs OXA and hycanthone to their active forms [23,24]. It displays sulfotransferase activity *in vitro* on exogenous substrates [23], even though the protein shows a low level of sequence similarity to other sulfotransferases, and it is mostly expressed in the intra-mammalian stages of the life cycle (schistosomula and adults, **Supplementary Figure S2A)** [25]. Intriguingly, *SULT-OR* belongs to a gene family that has expanded in trematodes [37], suggesting it may play an important role in clade-specific biology. However, *ex vivo SULT-OR* RNAi experiments in adult male worms showed no evident phenotypic effects other than becoming resistant to OXA [23]. Single-cell transcriptomic analysis of two-day-old schistosomula [9], adult worms [10], and *in vitro*-transformed mother sporocysts (unpublished) revealed *SULT-OR* mRNA is a marker of parenchymal cell clusters **(Supplementary Table S3** and **Supplementary Figure S2B)**, while its top BLASTP hit in the planarian *Schmidtea mediterranea*, *dd_Smed_v6_9472_0*, is a marker of intestinal cells [38].

### 3.2 A specific gRNA to introduce mutations in exon 1 of *SULT-OR*

The *SULT-OR* gene comprises two coding exons separated by one intron, spans 4837 bp on the reverse strand of chromosome 6, and includes a short 46 bp-5’ UTR **(Figure 1A)**. A gRNA was designed to target residues 19 to 38 of the coding region of *SULT-OR* within exon 1, adjacent to a TGG protospacer adjacent motif (PAM) and with the predicted double strand break (DSB; i.e. the predicted Cas9 cut site) 3 bp upstream of the PAM **(Figure 1B)**. Importantly, the sequences homologous to this gRNA’s target region are relatively diverged in the paralogous genes on chromosome 6, with many mismatches within the seed region (10-12 bp at its 5’ end) **(Supplementary Figure S3A)**. It has been shown that mismatches in the gRNA ‘core’ sequence located between 4 to 7 nucleotides upstream of the PAM abolishes off target cleavages [19, 39]; hence, our gRNA is expected to be specific to *SULT-OR*.

### 3.3 CRISPR-Cas9 machinery successfully delivered into schistosome developmental stages

To investigate whether the CRISPR-Cas9 machinery, i.e. RNP (ribonucleoprotein) complex containing the Cas9 nuclease and *SULT-OR*-specific gRNA, was successfully delivered into adult worms, sporocysts and eggs, we used fluorescently labelled RNP. Parasites were collected from culture four hours after transfection and fixed for confocal microscopy. The images revealed that the RNP complex entered cells of adult worms, sporocysts and eggs **(Figure 2)**. Even though the parasites were thoroughly washed before fixation, a strong signal outside the tegument was evident, in particular in adult worms, suggesting RNP complex molecules unspecifically bound to the surface of the parasites **(Figure 2A** and **Movie S1)**. However, in addition to the signal in the surface of the parasite, the confocal optical sections revealed fluorescently-labelled cells within the body of both male and female worms **(Figure 2B**, **Supplementary Figure S4A-C**, and **Movies S2** and **S3**). Interestingly, the majority of these successfully transfected cells were located around the intestine **(Figures 2B-D** and **Movie S1)**. The relatively higher concentration of the RNP complex surrounding the adult gut may have resulted from worms swallowing Cas9-gRNA molecules in the suspension before the electroporation step was carried out. The fluorescent signal in sporocysts was evenly distributed within the organism **(Figures 2E, G, Supplementary Figures S5A, B** and **Movie S4)**, whereas within eggs the signal was mainly localised outside the larvae, but inside the eggshell **(Figures 2F, H)**. Importantly, no autofluorescence signal was seen in control parasites **(Supplementary Figure S4D, F** and **Supplementary Figure S5C, D)**.

**Figure 2.**
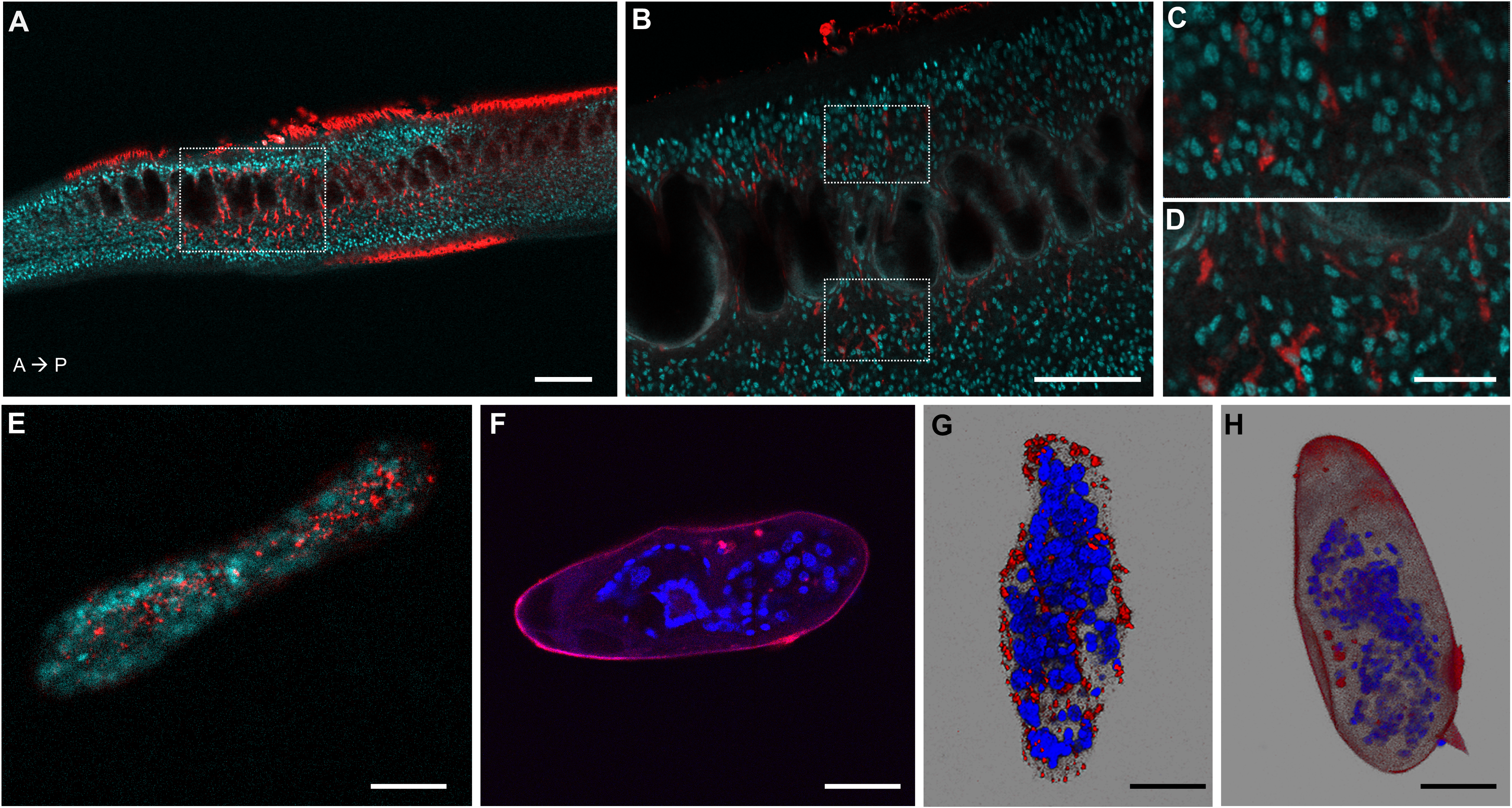
Confocal microscopy images of *S. mansoni* parasites transfected with fluorescently labelled Cas9-gRNA (ATTO™ 550 signal in red), fixed and DAPI-stained (DAPI signal in aqua blue or blue). **(A)** Confocal optical section of a male adult worm. A→ P: indicates the anterior-posterior axis. Scale bar: 100 μm. **(B)** Magnified squared-area in (A). Scale bar: 50 μm. **(C, D)** Magnified top and bottom squared-areas in (B), respectively. Scale bar: 10 μm. **(E, F)** Confocal optical sections of a sporocyst and an egg, respectively. **(G, H)** Maximum intensity projection of z-stack images of a sporocyst and an egg, respectively. Scale bars in E-H: 25 μm. The images of worms, sporocysts and eggs, were taken from representative specimens collected from the biological replicate 3, 2, and 1, respectively.

### 3.4 Evident CRISPR-Cas9-induced deletions in exon 1 of *SULT-OR*

The CRISPR-Cas9 transfection experiments were performed on adult worms (3 biological replicates), mother sporocysts (2 biological replicates) and eggs (3 biological replicates). All biological replicates were performed by different experimentalists on different days as indicated in **Supplementary Table S1**. DNA was extracted from the parasites four days after transfection, and a 262-bp PCR product spanning the gRNA region was amplified **(Figure 1C)** and sequenced on an Illumina MiSeq. The *SULT-OR-*specific PCR primers were designed in regions that are divergent between *SULT-OR* and the other paralogous genes **(Supplementary Figure S3B, C)**. With the assistance of the CRISPResso software [32, 33] we searched for mutations in the sequence reads by aligning reads to the reference amplicon. Remarkably, the percent of aligned reads that contained deletions was significantly higher for CRISPR-Cas9-treated samples than for matched controls when we pooled all tested developmental stages **(Figure 3A** and **Supplementary Table S1**; paired one-sided Wilcoxon test: *n*=8 biological replicates, *P*=0.04). In contrast, the percent of aligned reads with insertions or substitutions was not consistently higher in CRISPR-Cas9-treated samples than matched controls (*P*=0.2 for insertions and *P*=0.9 for substitutions; **Supplementary Figure S6B, C)**. The apparent substitutions seen in both CRISPR-treated and control samples are likely due to sequencing errors, especially at the ends of the amplicon, since the two reads of a read-pair overlap in a 38-bp region in the centre of the amplicon, allowing CRISPResso to infer a higher-quality consensus sequence for that central region **(Figure 1C)**.

**Figure 3.**
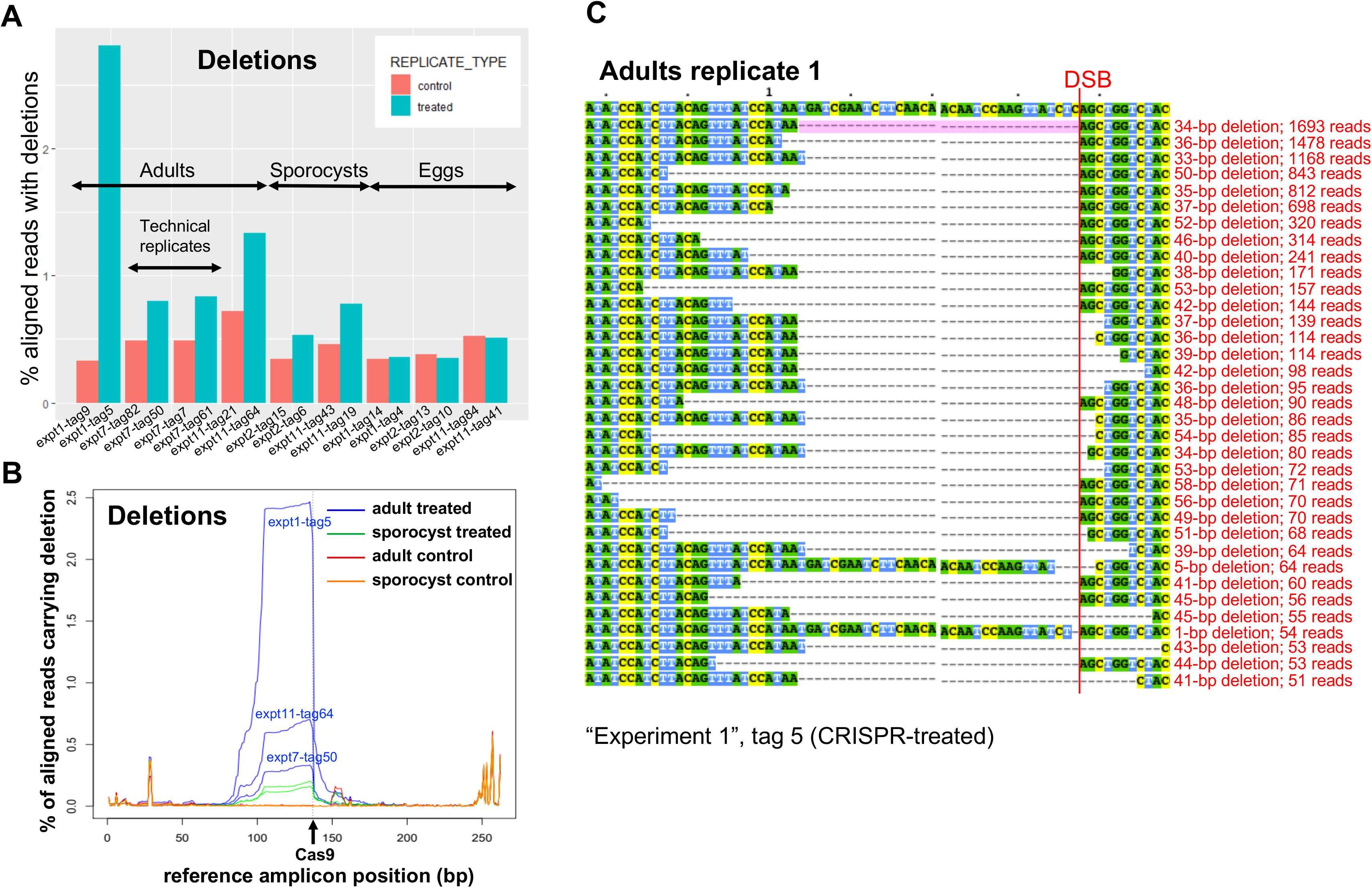
**(A)** Frequency of deletions in NGS sequencing data, identified with the assistance of CRISPResso in three biological replicates from adults, two from sporocysts, and three from eggs. **(B)** CRISPR-induced deletions in adult worms and sporocysts. The positions of deletions found by CRISPResso in the reference amplicon are indicated, in three biological replicates of CRISPR-Cas9-treated adult samples (blue lines: experiment 1, tag 5; experiment 7, tag 50; experiment 11, tag 64) and matched adult control samples (red lines: experiment 1, tag 9; experiment 7, tag 82; experiment 11, tag 21), and in two biological replicates of CRISPR-Cas9-treated sporocysts (green lines: experiment 2, tag 6; experiment 11, tag 19) and matched sporocyst controls (orange lines: experiment 2, tag 15; experiment 11, tag 43). The black arrow shows the predicted Cas9 cut site. **(C)** Multi-sequence alignment of *SULT-OR* alleles with deletions found in CRISPR-Cas9-treated adult worms that are supported by >=50 reads and span the DSB site indicated with a red line, based on one of the treated adult replicates (experiment 1, tag 5). The common 34-bp deletion is highlighted in pale pink.

Remarkably, a closer examination revealed that all three biological replicates of CRISPR-Cas9-treated adult worms had large deletions absent from control samples, extending from the predicted Cas9 cut site to about 60 bp upstream **(Figure 3B, C)**. Considering the reads that contained a single internal deletion spanning the predicted Cas9 cut site, and no internal insertions, we found that 0.3-2.0% of aligned reads from CRISPR-Cas9-treated adult worms exhibited such deletions, compared to 0.0% of aligned reads from matched controls **(Supplementary Table S1)**.

### 3.5 Higher CRISPR-induced mutation rate in adults compared to sporocysts and eggs

Interestingly, up to 10 times more reads containing deletions spanning the predicted Cas9 cut site were detected in CRISPR-Cas9-treated adult worms (0.3-2.0% of aligned reads) compared to CRISPR-Cas9-treated sporocysts (0.1-0.2%; **Supplementary Table S1)**. In contrast, in eggs the rate of such deletions was not any higher than in matched controls **(Supplementary Table S1)**.

Deletions of the same size, and in the same position, were identified in CRISPR-Cas9-treated sporocysts and adults, being absent from respective matched controls **(Figure 3B)**. In addition, across different biological replicates of CRISPR-Cas9-treated adults and sporocysts, the most frequent deletion alleles (i.e. those for which we detected the most supporting reads) had roughly the same sizes and positions **(Figure 3C** and **Supplementary Figure S7)**. The most common deletion identified in all three adult biological replicates, and in one of the two sporocyst biological replicates, was 34 bp directly upstream of the predicted Cas9 cut site (spanning positions 104-137 in the reference amplicon). We observed rare deletions that were up to three times longer: that is, deletions that extended from the predicted Cas9 cut site to 102 bp upstream (to position 36 in the reference amplicon). Strikingly, none of these deletions were apparent in CRISPR-Cas9-treated eggs.

Almost all the deletions observed extended upstream from the predicted Cas9 cut site; rare deletions extending both upstream and downstream of the cut site were identified but at relatively lower frequency, although often supported by 50 or more reads **(Supplementary Figure S7).** In all biological replicates from adults, we did observe extremely low-frequency deletions, supported by few reads (<50 reads, not shown in **Supplementary Figure S7)**, extending from the predicted Cas9 cut site to 102 bp upstream (position 36 in the reference amplicon), and deletions spanning the cut site that extended as far as 79 bp downstream of the cut site (position 216 in the amplicon).

The percent of aligned reads carrying deletions that spanned the predicted Cas9 cut site did not differ between CRISPR-Cas9-treated eggs and control eggs. This suggested that in eggs either CRISPR-Cas9 did not introduce mutations in *SULT-OR* or they had occurred at an extremely low level. The presence of a recognition site for the restriction enzyme PvuII overlapping the predicted Cas9 cut site **(Figure 1B)** allowed us to develop a protocol to enrich for mutant alleles. Any CRISPR-Cas9-induced deletions that extended upstream from the Cas9 cut site would remove this PvuII recognition site, so by digesting the DNA from treated parasites with PvuII, we expected to enrich for CRISPR-Cas9-induced deletions. In two out of three biological replicates of CRISPR-Cas9-treated egg samples, after PvuII treatment we were able to detect a slightly higher rate of deletions spanning the predicted Cas9 cut site, compared to in PvuII-treated control egg samples, i.e. an increase of at least 2-fold **(Supplementary Table S1)**, even though these deletions were still at very low frequency. This finding indicates that CRISPR-Cas was indeed active in eggs, although at very low levels.

### 3.6 Evidence for large deletions

Large CRISPR-induced deletions of >500 bp have been observed in the nematode *Strongyloides stercoralis [20].* In addition to the most common CRISPR-Cas9-induced deletions observed in *S. mansoni* that extended 34 bp upstream of the predicted Cas9 cut site **(Figure 3B)**, we did observe low-frequency deletions (supported by few reads) extending from the predicted Cas9 cut site to 102 bp upstream (to position 36 in the reference amplicon) **(Figure 4** and **Supplementary Figure S8)**. We simulated reads carrying deletions of every possible length, extending upstream from the predicted Cas9 cut site, that is, a read carrying a deletion of 1-bp upstream of the Cas9 cut site, a read carrying a deletion of 2-bp upstream of the Cas9 cut site, reads with deletions of 3-bp, 4-bp, 5-bp, and so on. Using our parameter settings, CRISPResso detected the simulated deletions of sizes ranging from 1-bp up to 104-bp upstream of the predicted Cas9 cut site, but could not detect simulated deletions extending 105 bp or further upstream of the Cas9 cut site. This is presumably because, by default, CRISPResso requires that a read aligns with at least 60% identity to the reference amplicon, thus, discarding reads carrying simulated deletions spanning 40% (i.e. 105 bp) or more of our 262-bp reference amplicon. Given that in our real data **(Figure 4** and **Supplementary Figure S8)**, we observed low-frequency deletions extending up to 102 bp upstream of the predicted Cas9 cut site, it is possible that in reality even longer deletions did also occur, but they would not have been detected, either because the reads were discarded by CRISPResso or one or both PCR primer regions were deleted.

**Figure 4.**
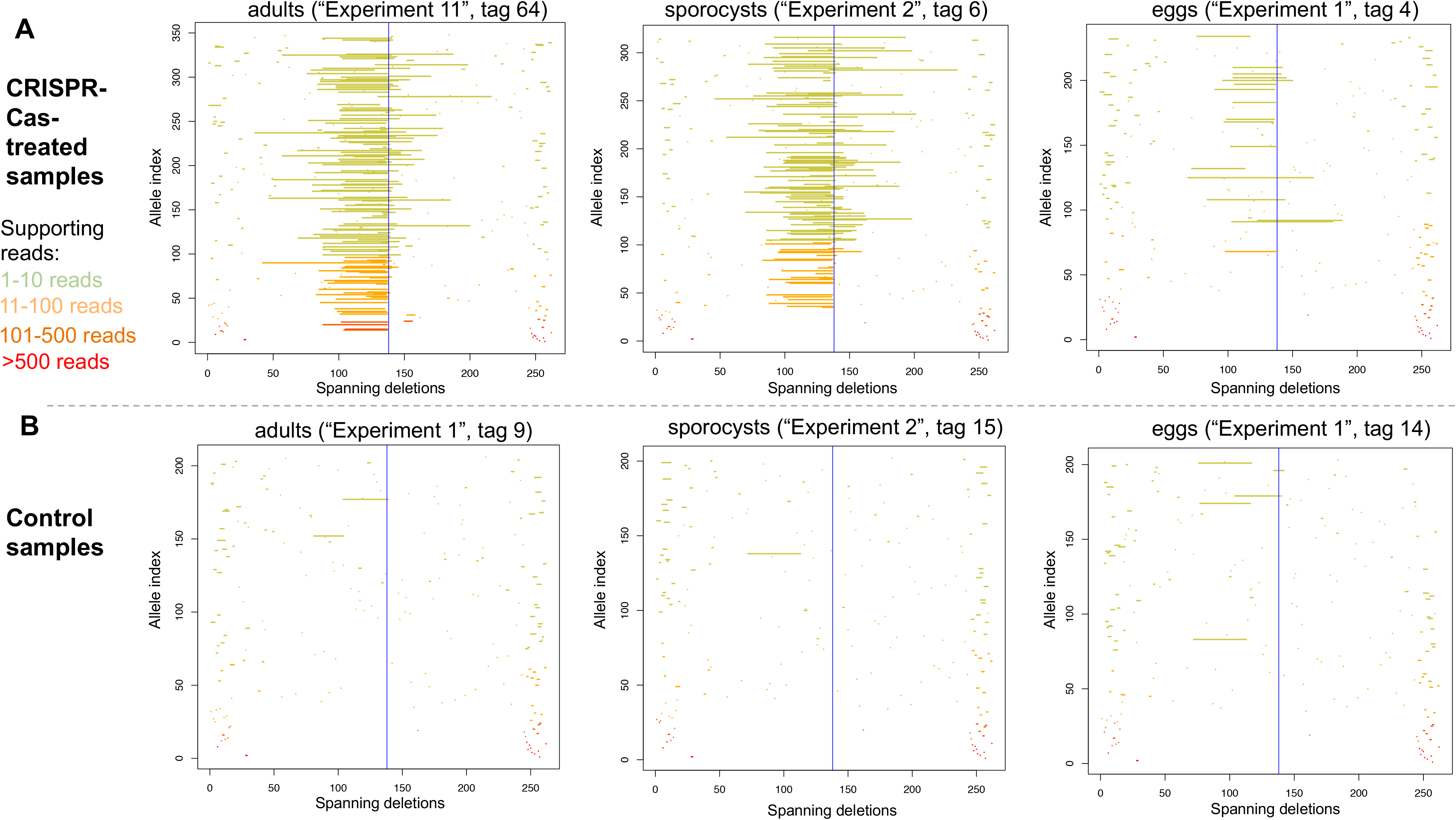
Deletion alleles seen in the *SULT-OR* gene in amplicon sequencing reads from treated **(A)** and control **(B)** adults (left), sporocysts (centre), and eggs (right), showing alleles that contain a single internal deletion and no internal insertions with respect to the reference amplicon. The y-axis shows deletion alleles sorted by the number of reads supporting them, with the alleles supported by the most reads at the bottom. Alleles supported by >500 reads in red, alleles supported by 101-500 reads in dark orange, alleles supported by 11-100 reads in pale orange, and alleles supported by 1-10 reads in pale green. The x-axis shows the position of the deletion along the reference amplicon, with a blue vertical line at the predicted Cas9 cut site.

### 3.7 Deletions predicted to cause oxamniquine resistance

Relative to the SULT-OR amplicon, the start codon is located at position 103 and the predicted Cas9 cut site at position 137. In adult worms and sporocysts, the CRISPR-Cas9-induced deletions extended upstream from the predicted cut site into the first exon. The most common deletions extended 34 bases upstream, completely removing the first coding exon (barring a single a base) and shifting the reading frame of the entire coding region **(Figure 1C** and **Supplementary Figure S7)**. This would result in the preferential degradation of the mutant mRNAs, as previously suggested to explain the *ω1* knockdown at the mRNA level observed in CRISPR-Cas9-treated eggs [21]. Moreover, any remaining frame-shifted SULT-OR protein is predicted to have lost its ability to convert OXA to its active drug form. Furthermore, even longer deletions upstream of the predicted Cas9 cut site were observed **(Supplementary Figure S7)**, which may extend into the *SULT-OR* promoter region given the 5’-UTR spans only 46 bp upstream of the first protein-coding codon, and hence may ablate the transcription of *SULT-OR.* The large CRISPR-Cas9-induced deletions, if they are homozygous, are predicted to cause resistance to OXA, either by producing frameshifts in the *SULT-OR* mRNA or by ablating *SULT-OR* transcription. However, when we analysed the expression levels of *SULT-OR* mRNA by qRTPCR across preparations of whole parasites in CRISPR-Cas9-treated samples versus control adult worms, no differences were evident (not shown).

## 4. Discussion

Genome editing mediated by CRISPR-Cas has been recently applied to *S. mansoni* to knock out the egg-specific gene *omega-1* (*ω1*) [21]. The CRISPR-Cas9 treatment of eggs by electroporation in the presence of the RNP complex, or egg transduction with lentivirus particles expressing Cas9 and the gRNA, induced a detectable knock-down both at the mRNA and protein levels and a clear phenotype of smaller granulomas in mice exposed to CRISPR-Cas9-treated eggs. In the current study, we decided to employ CRISPR-Cas9 to target the *SULT-OR* sulfotransferase gene in *S. mansoni* with an RNP complex. Strikingly, we detected large deletions of ≥34 bp extending upstream of the predicted Cas9 cut site, whereas deletions extending downstream of the cut site were extremely rare. The tendency for deletions to be upstream of the predicted Cas9 cut site agrees with observations in mouse cell lines [40]. We identified deletions extending up to 102 bp upstream of the predicted Cas9 cut site, reaching the limit detectable with CRISPResso (104 bp upstream, using our own parameter settings), suggesting that even larger deletions may have been missed. Deletions of several hundred base pairs have been described in *Strongyloides* [20], *C. elegans* [41], and mammalian cell-lines [40,42]. In addition, we characterised CRISPR-Cas9-induced mutations across three discrete developmental stages: adult worms, eggs, and the *in vitro*-transformed sporocysts. The deletions spanning the predicted Cas9 cut site were most commonly detected in adult worms (0.3-2.0% of aligned reads), followed by sporocysts (0.1-0.2%), and extremely rare in eggs. Interestingly, no evidence for CRISPR-Cas9-induced insertions or substitutions in the *SULT-OR* gene was observed. An interesting question is whether the pattern of CRISPR-Cas-induced mutations varies from gene-to-gene. When targeting the *ω1* gene in the absence of donor molecule, where CRISPR-Cas9-induced mutations presumably occurred by non-homologous end joining (NHEJ) [19], the overall rate of deletions detected in reads from CRISPR-Cas9-treated eggs was not higher than in control eggs [21]. This is consistent with an extremely low rate of NHEJ-mediated deletions in the *ω1* gene in eggs, similar to what we described here for the genome editing of the *SULT-OR* gene in eggs. On the other hand, when the *ω1* gene was targeted using CRISPR-Cas9 in the presence of a ssODN donor molecule, rare larger deletions spanning the predicted Cas9 cut site were detected in the CRISPR-Cas9-treated eggs compared to controls [21]. Since this effect was only observed in the presence of the donor molecule, it is tempting to speculate that those large deletions in the *ω1* gene may have been related to the homology directed repair (HDR) mechanism involved in the knock-in of the donor molecule rather than driven by NHEJ [43]. In addition, differences between protocols employed for *ω1* and *SULT-OR* may have resulted in different spectra of CRISPR-Cas9-induced mutations: the CRISPR-Cas9-treated eggs sequenced in the *ω1* study were transduced with lentivirus encoding a single-gRNA and Cas9 nuclease [21], whereas we induced mutations in *SULT-OR* by electroporating the parasites with an *in vitro*-assembled RNP complex of Cas9 nuclease and a two-piece gRNA. Interestingly, in the liver fluke *Opisthorchis viverrini* electroporated in the presence of a plasmid encoding a single-gRNA and Cas9 nuclease, small deletions of up to ~10 bp and insertions of up to 2 bp were detected near the predicted Cas9 cut site in the granulin gene [22].

Our data suggest that for *SULT-OR* the highest CRISPR efficiency was in adults, followed by sporocysts, and the lowest efficiency in eggs. For the latter we only found such deletions in <0.02% of aligned reads even after using PvuII to enrich for mutant reads. Three possible non-mutually exclusive hypotheses may explain these differences. Firstly, electroporation of the RNP complex may be most efficient in adults and least efficient in eggs, possibly because the surface area:volume ratio of an adult worm is greater than that of a sporocyst or egg, and/or because the egg has a protective coating that makes it hard to penetrate [44]. However, microporous and internal microcanals shown to be scattered across *Schistosoma* eggshells would allow the interchange of macromolecules with the host tissues [45]. Relevant for us, the diameter of the smallest pores in *S. mansoni* eggs is 100 nm, and we have estimated the diameter of the RNP complex, assuming a globular shape, to be ~10 nm [46] indicating that the complex could have entered the egg through the pores. A second possibility is that some key NHEJ repair enzymes required for CRISPR have higher expression in adults than in sporocysts or eggs; according to RNAseq metadata [25] this is the case for *Smp_211060*, previously identified [21] as a homolog of the Ku70/Ku80 genes that play a key role in NHEJ [47]. Finally, CRISPR might be more efficient in inducing mutations in *SULT-OR* in adults than sporocysts or eggs, because *SULT-OR* is expressed more in the former developmental stage, making its chromatin more open and therefore more accessible to the CRISPR machinery [19,48–50].

To create a CRISPR-Cas9-mediated mutant of any schistosome gene in every cell of the animal, the germline cells need to be targeted and mutated by a germline transgenesis approach. So far, only two studies demonstrated germline transmission of exogenous DNA in schistosomes. An early study published in 2007 [51] showed that a GFP-expressing plasmid was introduced into the miracidium germ cells by particle bombardment. Subsequently, the transfected miracidia infected snails, and resulting cercariae were employed to infect hamsters and obtain F1 transformed eggs. However, over a few generations the transformed parasites died and/or (as expected) the plasmid was diluted or lost. A few years later, germline transmission of integrated retroviral transgenes was demonstrated. Murine Leukemia Retrovirus (MLV) transgenes transduced the germ cells of eggs and were propagated through both the intrasnail asexual developmental stages and intramammalian sexual developmental stages, reaching the F1 eggs [16, 17]. However, no germline transgenesis approach has yet been achieved using CRISPR-Cas in schistosomes. Ittiprasert *et al* in 2019 [21] applied the CRISPR technology using a lentivirus expressing Cas9 and the gRNA, plus a donor, to introduce a 24-bp insertion (by HDR) into the *ω1* gene in *S. mansoni* eggs. Intriguingly, although the expression of the *ω1* gene was reduced by 81-83% after CRISPR-Cas9 treatment using the donor molecule, only ~4.5% of reads were identified by amplicon sequencing as mutated by indels (with <1% showing deletions) or substitutions, and only 0.19% of reads contained the 24-bp insertion. Proposed explanations for this discrepancy include a preferential penetration of CRISPR-Cas9 machinery (lentivirus and donor) in the envelope of the egg, where *ω1* may be expressed [44], and/or the presence of large deletions that removed either one or both primer regions and so were not detected by amplicon sequencing (as seen in *Strongyloides* [20]). Similarly, expression of the *Opisthorchis viverrini* granulin gene was reduced by >80% after CRISPR-Cas9 treatment of pooled adults in the absence of a donor, but only 1.3% of amplicon sequencing reads contained indel mutations [22]. This apparent anomaly may be due to the predominant expression of the granulin gene in the *O. viverrini* tegument and gut, where electroporation of the gene editing plasmid may have been most efficient [22]. Furthermore, there may be variation in CRISPR-Cas9 efficiency between individual adults: taking adult *Opisthorchis* worms from hamsters that had been infected 60 days previously with CRISPR-treated newly encysted juveniles (NEJs), individual adults in which there was a greater knock-down at the mRNA level showed a far greater level of mutations upon amplicon sequencing, especially deletions and substitutions [22]. In this species, significant variation was seen in CRISPR-Cas9 efficiency between life stages. Using a plasmid encoding Cas9 and gRNA, a knock-down of >80% of granulin mRNA levels was achieved in adults and NEJs, but of <4% in metacercariae, possibly due to inefficient electroporation of the plasmid through the metacercarial cyst wall [22].

In our study, while amplicon sequencing revealed reads carrying CRISPR-Cas9-induced mutations in *SULT-OR*, a knock-down of *SULT-OR* at the mRNA level was not evident, probably because the deletions occurred in only a small fraction of the adult cells that express *SULT-OR*. Single-cell sequencing data from adults shows *SULT-OR* is a marker of parenchymal cells [10], while our confocal microscopy data suggest the Cas9-gRNA complex penetrated better into the adult tegument and intestine compared to parenchymal tissue. The same phenomenon was previously described in the liver fluke *Fasciola hepatica* when delivering fluorescently labelled molecules by electroporation [52,53], suggesting the flatworm intestine as the main point of entry when this delivery approach is employed. Furthermore, *SULT-OR* is also expressed at a low level in many other cell types in adults **(Supplementary Figure S2B)**. The large deletions spanning the predicted Cas9 cut site were found in 0.3-2.0% of aligned reads from CRISPR-treated adult worms, so our best estimate of the fraction of adult cells in which CRISPR worked is 0.3-2.0%. Since a pool of five adult worms were transfected with the RNP complex, the efficiency of CRISPR (and the amount of knock-down at the mRNA level) may have varied between worms, as well as between cells of an individual worm.

To conclude, more work is required to optimise CRISPR-Cas protocols to work best at different developmental stages and in particular tissues, and understand whether these differing protocols will result in different spectra of mutations [41] and degrees of mRNA knock-down. To do this, it may be critical to identify the mechanisms underlying CRISPR-Cas-induced mutations in schistosomes in each case (e.g. NHEJ, HDR or other mechanisms such as polymerase theta-mediated end-joining [54]). Addressing these items would help the research community to achieve the holy grail of targeting the germ line and creating a stable knock-out or knock-in strain of any gene of interest.

## Acknowledgments

We are grateful to colleagues at the Wellcome Sanger Institute; Simon Clare, Cordelia Brandt, Catherine McCarthy, Katherine Harcourt and Lisa Seymour for assistance and technical support with animal infections and maintenance of the *Schistosoma mansoni* life cycle; Carmen L. Diaz for sharing unpublished scRNAseq data from *in vitro*-sporocysts; Kate Rawlinson, Claire Cormie and David Goulding for technical assistance with the confocal microscopy; Steve Doyle, Marcus Lee, Katharina Boroviak, Sophie Adjalley and Andrew Bassett for informative discussions. We would like to thank James J. Collins III from The University of Texas Southwestern Medical Center, USA, for sharing scRNAseq data from adult worms. This study was supported by the Wellcome Strategic Award number 107475/Z/15/Z. The Wellcome Trust provided core-funding support to the Wellcome Sanger Institute, award number 206194.

## Author Contribution

Conceptualization: MB, GR; Data Curation: AC, AT; Formal Analysis: GS, AC, PD, GR; Funding Acquisition: MB; Investigation: GS, AC, PD, ML, GR; Methodology: GS, GR; Project Administration: GR, NH, MS; Resources: MB, GR; Software: AC; Supervision: GR; Validation: GS, AC, PD, GR; Visualization: AC, GR; Writing – Original Draft Preparation: AC, GR; Writing – Review & Editing: all authors.

## Supplementary information

### Supplementary Figures

**Supplementary Figure S1. (A)** Phylogenetic tree of the Compara gene family that contains the *S. mansoni SULT-OR* gene (*Smp_089320*). *SULT-OR* belongs to a clade of paralogous genes that lie about 3.2 Mb along chromosome 6. This clade also contains haplotypic gene copies that lie on a chromosome 6 haplotypic contig (SM_V7_6H002). Adapted from WormBase ParaSite v14 [34]. **(B)** The region of chromosome 6 that contains the *SULT-OR* gene and paralogous genes belonging to the same clade (top), and the haplotypic contig SM_V7_6H002 containing haplotypic gene copies belonging to that clade (bottom). Note that the gene *Smp_315270* on SM_V7_6H002 is homologous to *SULT-OR* but belongs to a different Compara gene family in WormBase ParaSite.

**Supplementary Figure S2. (A)** Expression profile of *SULT-OR* across S*. mansoni* developmental stages. Gene expression in male and female parasites is indicated in blue and red, respectively. Adapted from https://v7test.schisto.xyz [25]. **(B)** Uniform Manifold Approximation projection (UMAP) plot of the *SULT-OR* gene (*Smp_089320*) expression pattern in adult worms showing an enriched expression in the parenchyma_1 (1), parenchyma_2 (2) and unknown (3) cell clusters. Raw data obtained from [10].

**Supplementary Figure S3. (A)** Multiple sequence alignment of the coding region of exon 1 of *SULT-OR* and related genes in the same clade of the sulfotransferase gene family. The gene *Smp_336300* is in the same clade but is quite diverged in its exon 1 with indels with respect to *SULT-OR*, and therefore was not included in the alignment. The start codon and position of the gRNA are indicated in red boxes. **(B)** Multiple sequence alignment of the 117 bp-upstream sequence from the ‘ATG’ start codon of *SULT-OR* (indicated in red box), and the same length of sequence upstream of related genes in the same clade, excluding *Smp_336300*. For *Smp_328800*, the likely start codon was taken to be the next in-frame ‘ATG’ downstream of the start codon given in WormBase ParaSite, based on comparison of its predicted protein-coding sequence to that of *SULT-OR*. The forward primer sequence used to generate the amplicon libraries is indicated in the red box. **(C)** Multiple sequence alignment of the coding region of exon 1 of *SULT-OR* downstream of the Y36 codon (indicated in the red box), with homologous regions of related genes in the same clade. The annealing sequence of the reverse primer used to generate the amplicon libraries is indicated in the red box. The alignments were made using Clustal-omega v1.2.4 [55], and viewed using Mview v1.63 [56].

**Supplementary Figure S4.** Confocal microscopy images of *S. mansoni* adult worms transfected with fluorescently labelled Cas9-gRNA (ATTO™ 550 signal in red), fixed and DAPI-stained (DAPI signal in aqua blue), and untreated controls. **(A)** Confocal optical section of a fluorescently labelled Cas9-gRNA-treated male worm. Scale bar:100 μm. **(B)** Magnified squared-area in (A). Scale bar: 25 μm. **(C)** Magnified squared-area in (B). Scale bar: 10 μm. **(D, E)** Confocal optical sections of an untreated control or a fluorescently labelled Cas9-gRNA-treated male worm, respectively. **(F, G)** Confocal optical sections of an untreated control or a fluorescently labelled Cas9-gRNA-treated female worm, respectively. Scale bars in D-G: 100 μm.

**Supplementary Figure S5.** Confocal microscopy images of *S. mansoni* sporocysts and eggs transfected with fluorescently labelled Cas9-gRNA (ATTO™ 550 signal in red), fixed and DAPI-stained (DAPI signal in aqua blue or blue), and untreated controls. **(A, B)** Confocal optical sections of fluorescently labelled Cas9-gRNA-treated sporocysts. **(C, D)** Confocal optical sections of an untreated control sporocyst and egg, respectively. Scale bars: 20 μm.

**Supplementary Figure S6.** Frequency of indels and substitutions in NGS sequencing data, identified using CRISPResso: **(A)** deletions, **(B)** insertions, and **(C)** substitutions, in the indicated samples.

**Supplementary Figure S7**. Deletion alleles found in CRISPR-Cas9-treated adult worms and sporocysts that span the predicted double stranded break site (DSB) and were supported by ≥ 50 reads. Reads containing just one internal deletion with respect to the reference amplicon, and no internal insertions were considered. Results for three different biological replicates from adults **(A-C)**, and for two biological replicates from sporocysts **(D, E)** are depicted. We did not see deletion alleles supported by >=50 reads and spanning the DSB site in matched control samples. The common 34-bp deletion is highlighted in pale pink. The alignments were displayed using Mview v1.63. [56].

**Supplementary Figure S8.** Deletion alleles seen in the *SULT-OR* gene in amplicon sequencing reads from additional replicates of CRISPR-treated adults, sporocysts, and eggs, as indicated, showing alleles that contain a single internal deletion and no internal insertions with respect to the reference amplicon. The y-axis shows deletion alleles sorted by the number of reads supporting them, with the alleles supported by the most reads at the bottom. Alleles supported by >500 reads in red, alleles supported by 101-500 reads in dark orange, alleles supported by 11-100 reads in pale orange, and alleles supported by 1-10 reads in pale green. The x-axis shows the position of the deletion along the reference amplicon, with a blue vertical line at the predicted Cas9 cut site.

### Supplementary Tables

**Supplementary Table S1.** Frequency of indels and substitutions in NGS sequencing data. Sample accession numbers are provided in columns P and Q.

**Supplementary Table S2.** Oligonucleotides used for dual-indexed sequencing library preparation and RT-qPCR oligonucleotides.

**Supplementary Table S3.** Expression profile of the *SULT-OR* gene in single-cell sequencing data. Raw single-cell sequencing data obtained from [9,10]

### Supplementary Movies

**Supplementary Movie S1.** Serial optical sections of a *S. mansoni* male adult worm transfected with fluorescently labelled Cas9-gRNA (ATTO™ 550 signal in red), fixed and DAPI-stained (DAPI signal in aqua blue). Scale bar: 100 μm.

**Supplementary Movie S2.** Serial optical sections of a *S. mansoni* male adult worm transfected with fluorescently labelled Cas9-gRNA (ATTO™ 550 signal in red), fixed and DAPI-stained (DAPI signal in aqua blue). In these series of optical sections the anterior end of the worm is observed. Scale bar: 100 μm.

**Supplementary Movie S3.** Serial optical sections of a *S. mansoni* female adult worm transfected with fluorescently labelled Cas9-gRNA (ATTO™ 550 signal in red), fixed and DAPI-stained (DAPI signal in aqua blue). In these series of optical sections the anterior end of the worm is observed. Scale bar: 100 μm.

**Supplementary Movie S4.** Serial optical sections of a *S. mansoni* sporocyst transfected with fluorescently labelled Cas9-gRNA (ATTO™ 550 signal in red), fixed and DAPI-stained (DAPI signal in aqua blue). Scale bar: 25 μm.

